# Evaluating the Genome and Resistome of Extensively Drug-Resistant *Klebsiella pneumoniae* using Native DNA and RNA Nanopore Sequencing

**DOI:** 10.1101/482661

**Authors:** Miranda E. Pitt, Son H. Nguyen, Tânia P.S. Duarte, Mark A.T. Blaskovich, Matthew A. Cooper, Lachlan J.M. Coin

## Abstract

*Klebsiella pneumoniae* frequently harbour multidrug resistance and current methodologies are struggling to rapidly discern feasible antibiotics to treat these infections. While rapid DNA sequencing has been proposed for prediction of resistance profile; the role of rapid RNA sequencing has yet to be fully explored. The MinION sequencer can sequence native DNA and RNA in real-time, providing an opportunity to contrast the utility of DNA and RNA for prediction of drug susceptibility. This study interrogated the genome and transcriptome of four extensively drug-resistant (XDR) *K. pneumoniae* clinical isolates. The majority of acquired resistance (≥75%) resided on plasmids including several megaplasmids (≥100 kbp). DNA sequencing identified most resistance genes (≥70%) within 2 hours of sequencing. Direct RNA sequencing (with a ∼6x slower pore translocation) was able to identify ≥35% of resistance genes, including aminoglycoside, β-lactam, trimethoprim and sulphonamide and also quinolone, rifampicin, fosfomycin and phenicol in some isolates, within 10 hours of sequencing. Polymyxin-resistant isolates showed a heightened transcription of *phoPQ (*≥2-fold) and the *pmrHFIJKLM* operon (≥8-fold). Expression levels estimated from direct RNA sequencing displayed strong correlation (Pearson: 0.86) compared to qRT-PCR across 11 resistance genes. Overall, MinION sequencing rapidly detected the XDR *K. pneumoniae* resistome and direct RNA sequencing revealed differential expression of these genes.

## INTRODUCTION

*Klebsiella pneumoniae* is one of the leading causes of nosocomial infections, with reports of mortality rates as high as 50% (1-5). This opportunistic pathogen frequently exhibits multidrug resistance which severely limits treatment options (6). A high abundance of resistance is commonly encoded on plasmids, accounting for the rapid global dissemination of resistance (1,6). Common therapeutic options for multidrug-resistant infections include carbapenems, fosfomycin, tigecycline and polymyxins (7). However, resistance is also rapidly developing against these antibiotics (6). Alarmingly, pandrug-resistant (PDR) *K. pneumoniae* have emerged which are resistant to all commercially available antibiotics (8,9).

One of the major contributors to the advent of antibiotic resistance is the inability for current detection methodologies to readily and accurately assess bacterial infections in particular, the resistance profile (10). This has resulted in the unnecessary use of antibiotics for viral infections and ineffective antibiotics being administered for resistant infections. Rapid sequencing has been proposed as a way to determine pandrug resistance profiles, including approaches which utilise high accuracy short reads, as well as those which exploit real-time single-molecule sequencing such as Oxford Nanopore Technologies (ONT). The ONT MinION platform is a portable single-molecule sequencer which can sequence long fragments of DNA and stream the sequence data for further data processing in real-time, detecting the presence of bacterial species and acquired resistance genes (11-15). Moreover, the long reads coupled with the ability to multiplex samples has immensely aided with the assembly of bacterial genomes (16-18). This capability allows for the rapid determination of whether resistance is residing on the chromosome or plasmid/s. Of particular interest are high levels of resistance encoded on plasmids, as these genes can rapidly be transferred throughout the bacterial population via horizontal gene transfer.

ONT has recently released a direct RNA sequencing capability, which sequences native transcripts. Other sequencing technologies rely on fragmentation, cDNA conversion and PCR steps which create experimental bias and hinder the accuracy of determining gene expression (19,20). The ability for MinION sequencing to read long fragments enables full length transcripts to be investigated. To date, only a few direct RNA sequencing publications exist which include eukaryote transcriptomes, primarily yeast (*Saccharomyces cerevisiae (*19,21)) and recently, human (BioRxiv: https://doi.org/10.1101/459529). This sequencing has additionally been implemented in viral transcriptomics (22, BioRxiv: https://doi.org/10.1101/300384, BioRxiv: https://doi.org/10.1101/373522). Only one prior study by Smith AM *et al.* has applied this sequencing to bacterial 16S ribosomal RNA (rRNA) to detect epigenetic modifications (BioRxiv: https://doi.org/10.1101/132274). Bacterial transcription differs significantly from eukaryotes in that transcription and translation occur simultaneously. As a result, bacterial mRNA transcripts lack poly(A) tails and alternative splicing (23). The poly(A) tail is critical for the library preparation for ONT sequencing thus, we have established a methodology for adding this component onto transcripts.

In this study, we applied MinION sequencing to interrogate both the genome and the transcriptome (via direct RNA sequencing) for XDR *K. pneumoniae* clinical isolates. Of interest was to compare the potential for RNA sequencing to provide a better correlation to the resistance phenotype than DNA sequencing. These isolates have previously undergone ‘traditional’ whole genome sequencing (Illumina) and antimicrobial susceptibility testing (24). Three strains were selected from this cohort which exhibited resistance to all 24 classes or combinations of antibiotics tested, a high abundance of antibiotic resistance genes (≥26) and differing lineages (ST11 (16_GR_13), ST147 (1_GR_13) and ST258 (2_GR_12)). Additionally, these isolates harbour polymyxin resistance which is facilitated by a disruption in or upstream of *mgrB*. MgrB is the negative regulator of PhoPQ and mutation results in the up-regulation of *pmrC* and the *pmrHFIJKLM* operon (25-27). This enables the addition of phosphoethanolamine and/ or 4-amino-4-deoxy-L-arabinose (Ara4N) onto the basal component of lipopolysaccharide, lipid A. These modifications perturb the key electrostatic interaction between lipid A and polymyxins that is critical for their activity (28,29). These pathways associated with polymyxin resistance were further explored using direct RNA sequencing. An additional polymyxin-susceptible XDR isolate (ST258; 20_GR_12) was selected to determine the differential expression associated with polymyxin resistance. This research aimed to assemble these genomes, discern the differential expression of resistance genes and ascertain the time required for detection. Furthermore, we sought to compare DNA and RNA sequencing as modalities for the rapid identification of acquired antibiotic resistance.

## MATERIAL AND METHODS

### Bacterial strains and growth conditions

Clinically acquired XDR *K. pneumoniae* strains were sourced through the Hygeia General Hospital, Athens, Greece (24). Antimicrobial susceptibility assays (Supplementary Table S1), sequence typing and detection of acquired resistance genes for these isolates have previously been determined (24). Strains were stored at −80°C in 20% (v/v) glycerol and the same stock was used as per the prior study (24). When required for extractions, glycerol stocks were grown on lysogeny broth (LB) agar plates and 6 morphologically similar colonies were selected for inoculation. The inoculum was grown in LB overnight at 37°C shaking at 220 rpm. This overnight inoculum was used for both DNA and RNA extractions.

### DNA extraction and high molecular weight DNA isolation

DNA was extracted from 10 ml of overnight culture using the DNeasy Blood and Tissue Kit (Qiagen) according to manufacturer’s guidelines, with the addition of an enzymatic lysis buffer pre-treatment (60 mg/ml lysozyme). High molecular weight (HMW) DNA from the prior extraction was selected using the MagAttract HMW DNA Kit (Qiagen) as per manufacturer’s instructions. Subtle changes included a further proteinase K treatment on the DNA extracts at 56°C for 10 min followed by supplementation of RNase A (1 mg) for 15 min at room temperature. Several attempts at direct DNA extraction from bacterial cells were undertaken using the MagAttract HMW DNA kit, however, were unsuccessful with these isolates. Due to several issues with potential carbohydrate contamination (260/230 ratio: ≤0.3), 2_GR_12 was also purified with the Monarch^®^ PCR & DNA Cleanup Kit (New England BioLabs) using the protocol to isolate fragments >2000 bp. DNA and RNA contamination was quantitated using Qubit®2.0 (Thermo Fisher Scientific) and purity determined with a NanoDrop 1000 Spectrophotometer (Thermo Fisher Scientific). DNA fragment sizes were determined using the Genomic DNA ScreenTape & Reagents (Agilent) and sizes from 200 to >60000 bp were analyzed on a 4200 TapeStation System (Agilent) (Supplementary Figure S1).

### RNA extraction, mRNA enrichment and poly(A) ligation

The overnight culture was sub-cultured in 10 ml of cation-adjusted Muller Hinton Broth (caMHB) to reflect conditions used for minimum inhibitory concentration (MIC) assays. Cultures were grown to mid-log phase (OD_600_ = 0.5-0.6). RNA was extracted via the PureLink^™^ RNA Mini Kit (Thermo Fisher Scientific) as per manufacturer’s protocols which included using Homogenizer columns (Thermo Fisher Scientific). To remove DNA contamination, the TURBO DNA-free^™^ kit was implemented. A minor adjustment was an increased concentration of TURBO DNase (4 U) incubated at 37°C for 30 min. The RNeasy Mini Kit (Qiagen) clean up protocol was additionally used to purify and concentrate RNA samples. Ribosomal RNA was depleted via the MICROB*Express*^™^ Bacterial mRNA Enrichment Kit (Thermo Fisher Scientific). Minor protocol changes included adding ≥2 µg of DNA depleted RNA and the enriched mRNA was precipitated for 3 h at −20°C. Poly(A) ligation was performed using the Poly(A) Polymerase Tailing Kit (Astral Scientific) as per the manufacturer’s alternative protocol (4 U input of Poly(A) Polymerase). The input RNA concentration was ≥800 ng and RNA samples were incubated at 37°C for 1 h. Poly(A) ligated RNA was purified using Agencourt AmpureXP (Beckman Coulter Australia) beads (1:1 ratio). RNA and DNA contamination was quantitated using the Qubit®2.0 (ThermoFisher Scientific) and purity determined with a NanoDrop 1000 Spectrophotometer (Thermo Fisher Scientific). RNA fragment size was checked using an Agilent RNA 6000 Pico kit and run on a 2100 Bioanalyzer (Agilent Technologies) for the initial RNA extract, post ribosomal RNA depletion and after poly(A) ligation (Supplementary Figure S2).

### RNA extraction, mRNA enrichment and poly(A) ligation

RNA libraries (≥600 ng poly(A) ligated RNA) were prepared using the Direct RNA Sequencing kit (SQK-RNA001). The Rapid Barcoding Sequencing kit (SQK-RBK001) was used for HMW DNA samples (1_GR_13, 16_GR_13, 20_GR_12; 300 ng input each). Isolate 2_GR_12 (300 ng input) was prepared separately using the Rapid Sequencing Kit (SQK-RAD003). Libraries were sequenced with MinION R9.4 flowcells and the raw data (fast5 files) were base-called using Albacore 2.1.1. RNA reads were additionally base-called with Albacore 2.2.7.

### Real-time resistome detection emulation

The real-time emulation was performed post sequencing and the time required to detect antibiotic resistance was determined as previously described (14). Briefly, this pipeline aligns Albacore base-called reads via BWA-MEM (ArXiv: https://arxiv.org/abs/1303.3997) to an antibiotic resistance gene database. Antibiotic resistance genes were obtained from the ResFinder 3.0 database (30). This dataset comprises of 2131 genes which were clustered based on 90% identity to form 611 groups or gene families. The detection of false positives is reduced using the multiple sequence alignment software kalign2 (31), a probabilistic Finite State Machine (32) and once the alignment score reached a threshold, the resistance gene was reported.

### Assembly of genomes

To assemble genomes with both Illumina and ONT reads, SPAdes v3.10.1 (33) were utilised. Hybrid assemblers included npScarf (34) and Unicycler v0.3.1 (35). Assemblers using only ONT reads included Canu v1.5 (excluding reads <500bp) (36) and the combination of Minimap2 v2.1-r311 and Miniasm v0.2-r168-dirty; Racon (git commit 834442) were used in both cases to polish the assemblies (37,38). Consensus sequences were determined using Mauve (snapshot_2015-02-13) to construct the final assembly (39). The output from each assembly software is reported in Supplementary Table S2. Genomes were annotated using the Rapid Annotation using Subsystem Technology (RAST) which also provided a list of virulence genes (40). The location of acquired antibiotic resistance genes were determined using ResFinder 3.0 (30) and plasmids were identified via PlasmidFinder 1.3 (41). To discern if plasmid sequences have previously been reported, contigs underwent a BLASTn analysis against the National Center for Biotechnology Information (NCBI) database (https://blast.ncbi.nlm.nih.gov/Blast.cgi).

### RNA alignment and expression profiling

Base-called RNA reads were converted to DNA (uracil bases changed to thymine) and aligned using BWA-MEM (parameters: -k 11 -W20 -r10 -A1 -B1 -O1 -E1 -L0 –Y) to the updated genome assemblies. Due to the lack of introns and full length transcripts being obtained, BEDTools coverage (42) was used to ascertain the relative expression of resistance genes. This was normalized to the number of counts obtained for the housekeeping gene, *rpsL (*43), to compare against qRT-PCR results. Read alignments were further visualised using Integrative genomics viewer (IGV) 2.3.59 (44).

### Whole transcriptome differential gene expression

To identify genes which were differentially expressed between a pair of samples (x and y), we used a beta-binomial distribution to calculate the probability of observing less than or equal to x_g reads mapping to gene g in sample x, conditional on the total number of reads mapping to all genes (sum_g(x_g)), the number of reads in sample y mapping to gene g (y_g) as well as the total number of reads mapping to all genes in sample y (sum_g (y_g). This was calculated in R using the pbetabinom.ab function in the VGAM package, with q = x_g, size = sum_g’(x_g’), alpha = y_g +1; beta = sum_g’(y_g’) - y(g) +1. Genes for which this probability was less than a predefined threshold were deemed to be significantly under expressed in sample x given sample y. A similar statistic was used to check for over-expression.

### Quantitative real-time reverse transcriptase PCR (qRT-PCR)

First strand synthesis to generate cDNA (1 µg total DNase-depleted RNA) was performed using SuperScript III (Thermo Fisher Scientific) which was also used for MinION direct RNA sequencing library preparations. Primers used are displayed in Supplementary Table S4. Samples were prepared in triplicate via the SYBR Select Master Mix (Thermo Fisher Scientific) and expression detected using a ViiA 7 Real-time PCR system (Thermo Fisher Scientific). Cycling conditions include: Hold 50°C (2 min), 95°C (2 min) followed by 50 cycles of: 95°C (15 sec), 55°C (1 min). A melt curve was included to determine the specificity of the amplification and a no template control to detect contamination or primer dimers. Results were analysed with QuantStudioTM Real-Time PCR Software, triplicates were averaged, normalised to the housekeeping gene *rpsL* and relative expression determined via the 2^*-δδCT*^ method (45).

### Data availability

Whole genome sequencing of the 4 clinical isolates, including the recent assembly, has been deposited under BioProject PRJNA307517 (www.ncbi.nlm.nih.gov/bioproject/PRJNA307517). ONT DNA sequencing data has been deposited on the Sequence Read Archive (www.ncbi.nlm.nih.gov/sra/) under study SRP133040. Accession numbers are as follows: 1_GR_13 (SRR6747887), 2_GR_12 (SRR6747886), 16_GR_13 (SRR6747885) and 20_GR_12 (SRR6747884). ONT direct RNA sequencing data (pass and fail reads) have been deposited on the Sequence Read Archive (www.ncbi.nlm.nih.gov/sra/) under study SRP133040. Accession numbers are as follows: 1_GR_13 (SRR7719054), 2_GR_12 (SRR7719055), 16_GR_13 (SRR7719052) and 20_GR_12 (SRR7719053).

## RESULTS

### Discerning the location of acquired resistance in the genome

Utilising the capacity for MinION sequencing to read long fragments of DNA, the location of antibiotic resistance genes were clearly resolved (Table 1). All genomes were circular with the exception of 2_GR_12 where 3 plasmids remained linear. This was partly due to difficulties extracting DNA and not retaining long fragments (Supplementary Figure S1). Amongst the four isolates, the chromosome size ranged between 5.1-5.5 Mb which encoded resistance genes *blaSHV-11, fosA* and *oqxAB*. The majority of resistance (≥75%) mapped to plasmids.

**Table 1.**
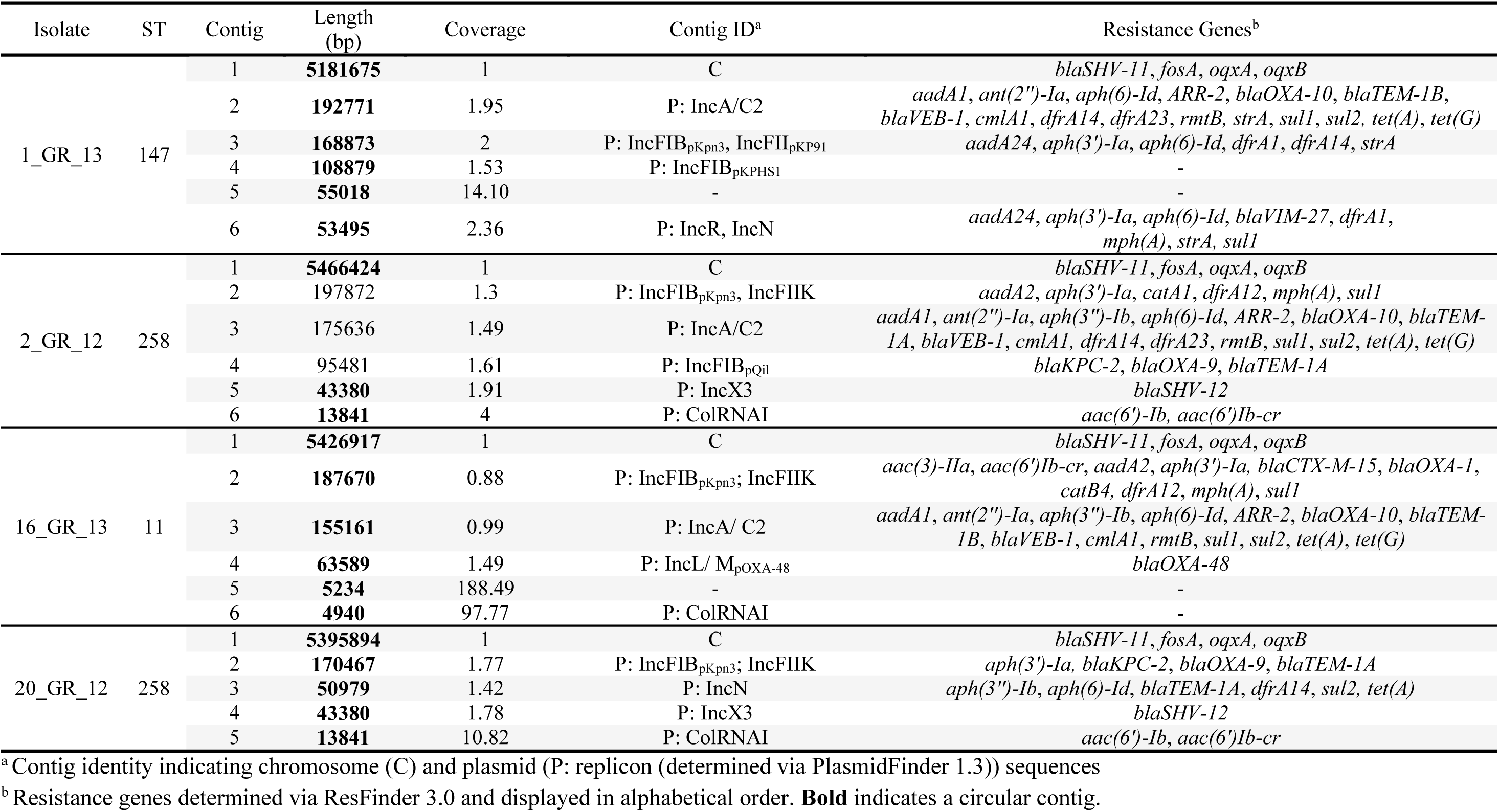
Final assembly of XDR *K. pneumoniae* isolates and location of antibiotic resistance genes

At least one megaplasmid, defined as a plasmid larger than 100 kbp, was detected in all isolates (Table 1). These commonly harboured the replicon IncA/C2 or InFIB and IncFIIK. The IncA/C2 plasmid was present in all samples except 20_GR_12. This plasmid contained up to 16 resistance genes which conferred resistance towards aminoglycosides, β-lactams, phenicols, rifampicin, sulphonamides, tetracyclines and trimethoprim, with the exception of 16_GR_13. Isolate 16_GR_13 lacked trimethoprim resistance on its IncA/C2 plasmid. The plasmids containing both replicons IncFIB and IncFIIK differed vastly between all four replicates. All contained IncFIB_pKpn3_ and IncFIIK, however, 1_GR_13 differed with IncFII_pKP91_. Additionally, a differing IncFIB replicon was detected on a separate contig in 1_GR_13 (pKPHS1) and 2_GR_12 (pQil). The only instance where another dual replicon was identified was in 1_GR_13 which harboured both IncR and IncN. This plasmid contained aminoglycoside, β-lactam, trimethoprim, macrolide and sulphonamide resistance. 1_GR_13 also contained a 5.5 kb circular contig which was annotated as a phage genome. Various regions of these megaplasmids were unique to these isolates compared to prior sequences deposited on NCBI (Supplementary Table S5).

The ColRNAI plasmid was present in all except 1_GR_13 which encoded aminoglycoside and quinolone resistance (*aac(6’)-Ib, aac(6’)-Ib-cr*) (Table 1). The ColRNAI plasmid in 2_GR_12 and 20_GR_12 was 13841 bp in size and shared 75% similarity between the two isolates. This plasmid differed in 16_GR_13 which contained no resistance genes and 35% the size. The same IncX3 plasmid (43380 bp) was apparent in isolates 2_GR_12 and 20_GR_12. Unique to 16_GR_13 was the IncL/ MpOXA-48 plasmid containing *blaOXA-48* and the 50979 bp IncN plasmid in 20_GR_12 with resistance against 5 classes (aminoglycoside (*aph(3’’)-Ib, aph(6)-Id*), β-lactam (*blaTEM-1A*), sulphonamide (*sul2*), tetracycline (*tet(A)*), trimethoprim (*dfrA14*)) of antibiotics.

Multiple copies of acquired resistance genes were apparent across plasmids in several isolates. For 1_GR_13, up to three copies were present of genes *aadA24, aph(3’)-Ia, aph(6)-Id, dfrA1, dfrA14, strA* and *sul1 (*Table 1). In 2_GR_12, *sul1* and *blaTEM-1A* were duplicated and for 16_GR_13, only *sul1* was represented twice.

### Real-time detection emulation of resistance genes via DNA sequencing

The vast majority (≥70%) of resistance genes were detected via DNA sequencing within the first 2 hours (Figure 1, Supplementary Table S3). These genes confer resistance towards aminoglycosides, β-lactams, fosfomycin, macrolides, phenicols, quinolones, rifampicin, sulphonamides, tetracyclines and trimethoprim. 20_GR_12 lacked acquired resistance genes for macrolides, phenicols and rifampicin, however, all other classes were detected within 2 hours. All isolates, except 2_GR_12, were sequenced for 21 hours which was sufficient to obtain the complete genome assembly. Only a few additional genes were detected after the first 10 hours across isolates (Supplementary Table S3). For 2_GR_12, an extended run of 41 hours detected no further genes after 20 hours. Overall, the presence of these resistance genes corresponded to a resistant phenotype towards aminoglycosides, β-lactams, fosfomycin, phenicols, quinolones, sulphonamides (sulfamethoxazole), tetracyclines and trimethoprim (Supplementary Table S1). As macrolides and rifampicin are not routinely used to treat *K. pneumoniae* infections, no breakpoints exist according to CLSI and EUCAST guidelines, hence, MICs were not determined.

**Figure 1.**
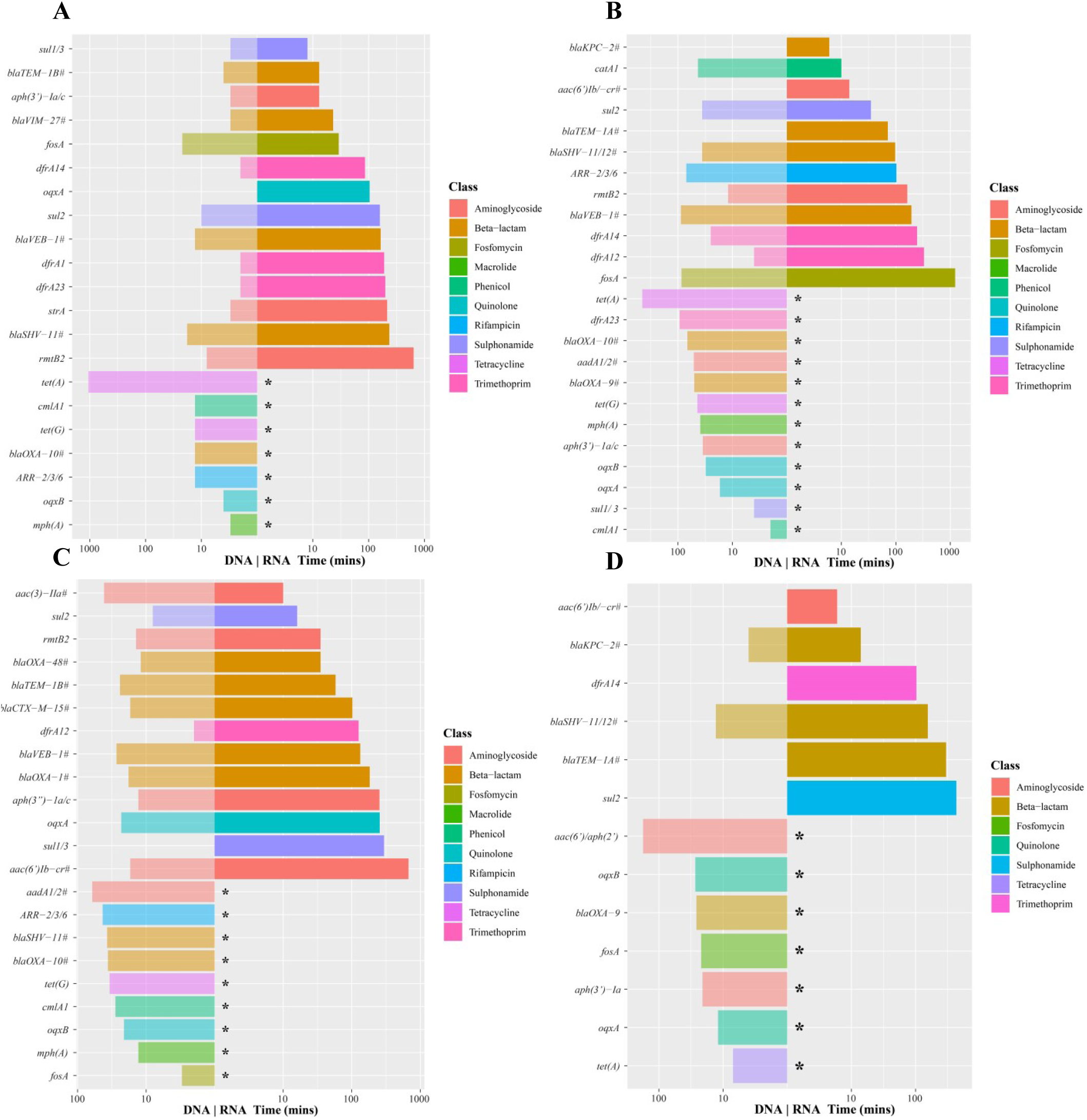
Time required to detect antibiotic resistance genes via the real-time emulation analysis using MinION DNA sequencing and direct RNA sequencing data. (**A**) 1_GR_13, (**B**) 2_GR_12, (**C**) 16_GR_13 and (**D**) 20_GR_12. Legend colours identify the class of antibiotic to which the gene confers resistance, / on y-axis indicates reads detected more than one resistance gene and # is a family of genes detected (>3). An asterisk (*) indicates the inability for direct RNA sequencing to detect this gene. Albacore 2.2.7 base-called sequences were used and all reads (pass and fail) were included in this analysis.

Post 2 hours of sequencing, several genes not observed in the final assembly via ResFinder 3.0 were detected (Supplementary Table S3). These were predominantly genes attributed to aminoglycoside, β-lactam, rifampicin and phenicol resistance. Furthermore, resistance genes to additional differing classes were detected including fusidic acid and vancomycin. This was evident in 2_GR_12 (*fusB*) and 16_GR_13 (*fusB, vanR*). However, these genes had less than 30 reads and their phred-scale mapping quality scores (MAPQ) were less than 10 (misplaced probability greater than 0.1). Furthermore, the majority of genes not observed in the final assembly nor observed in Illumina data exhibited a MAPQ score of ≤10 which may indicate that a more stringent threshold is required to negate false positives. However if this threshold increases, true positives would not be detected including *aadA1, aadA2* and *ARR-2* in 2_GR_12 and *blaOXA-48, blaCTX-M-15* and *ARR-2* in 16_GR_13.

Several genes found in the final assembly were not detected in the real-time emulation analysis (Supplementary Table S3). This was mainly observed for aminoglycoside resistance encoding genes. For 1_GR_13, this included *aadA1, ant(2’’)-Ia, aph(6)-Id* and *aadA24.* Similarly, 2_GR_12 and 20_GR_12 lacked *aph(3’’)-Ib* and *aph(6)-Id.* 2_GR_12 additionally had the absence of *ant(2’’)-Ia.* Detection of *ant(2’’)-Ia, aph(3’’)-Ib, aph(6)-Id* was not present in 16_GR_13. 16_GR_13 further lacked *catB4 (*phenicol) and *tet(A) (*tetracycline). Various phenicol resistance genes were reported in the real-time emulation however, the incorrect gene was identified which may represent sequencing errors accumulated over time and high similarity to other phenicol resistance genes. The tetracycline resistance gene, *tet(A)*, was interestingly not reported in this emulation with 190 reads and the majority of reads exhibiting a high mapping confidence (MAPQ = 60, equivalent to an error probability of 1×10^-6^). This gene was only detected after 10 hours for 1_GR_13 and 2_GR_12 and this result may be influenced by the presence of only 1 copy of *tet(A)* encoded on a low copy number megaplasmid (between 1 to 1.5, see Table 1).

### Direct RNA sequencing resistance detection

The time required to detect resistance was further interrogated using RNA sequencing. Rapid detection was apparent for several resistance genes via direct RNA sequencing (Figure 1). This was evident for genes conferring resistance to aminoglycosides, β-lactams, sulphonamides and trimethoprim for all four isolates. Resistance towards these antibiotics was commonly detected within 6 hours. In some instances, quinolone, rifampicin, fosfomycin and phenicol resistance was detected. This result remained similar when all reads or passed reads alone were analysed. The most significant difference when analysing all reads was the detection of *fosA* in 1_GR_13 and *ARR-2* and *fosA* in 2_GR_12. Consistently absent from this analysis were genes attributed to macrolide (*mph(A)*) and tetracycline (*tet(A), tet(G)*) resistance, however, isolates exhibited high levels of resistance to tetracycline (>64 µg/ml) (Supplementary Table S1). This may indicate that isolates require antibiotic exposure to enable transcription of these genes. Commonly no new genes were detected after 12 hours of sequencing with the exception of *fosA* in 2_GR_12. Although *fosA* was detected when including the failed reads, a low MAPQ score (≤10) was apparent. Similar to the DNA real-time detection, several genes not found in the final assembly were identified (Supplementary Table S3). With the exception of 20_GR_12, this included *aadB* and *strB* for all isolates. Additional genes detected included *ARR-7* in 1_GR_13, *strA* in 2_GR_12 and for 16_GR_13, *blaCTX-M-64, blaOXA-436* and *strA.* Similar genes or gene families were identified when comparing DNA and direct RNA sequencing. Overall, genes were detected more readily via DNA rather than RNA sequencing, possibly due to a lack of RNA expression in the absence of the antibiotic to which resistance is encoded. There were only a few instances where RNA sequencing detected resistance more quickly than DNA sequencing which included *aac(3’)-IIa* in 16_GR_13 and *sul2* and *catA1* in 2_GR_12. Similar results were apparent when investigating data yield rather than time (Supplementary Figure S4).

### Levels of expression of resistance genes

RNA sequencing accumulated over approximately 40 hours yielded between 0.9 and 1.7 million reads for these isolates (Supplementary Figure S3). However, only 10 to 14% of these reads successfully passed base-calling which was similar when using either Albacore 2.1.1 or 2.2.7. The low proportion of base-called reads reflects the fact that base-calling algorithms have not yet been optimised for direct RNA sequencing, and even less so for bacterial RNA sequencing. When aligning passed reads to the final assembly, ≥98% of reads were mappable, however, ≤40% of these had a MAPQ score ≥10. When all reads were aligned, ≤22% mapped to the genome and ≤5% exhibited a MAPQ score ≥10. A proportion of these reads were found to map to rRNA including 1_GR_13 (18%), 2_GR_12 (37%), 16_GR_13 (24%) and 20_GR_12 (23%). Overall, at least 58% of genes (with at least 1 read mapping to the gene) were identified to be expressed across isolates (1_GR_13 (68%), 2_GR_12 (58%), 16_GR_13 (75%) and 20_GR_12 (69%).

Amongst the four isolates, levels of expression for resistance genes on the chromosome (*blaSHV-11, fosA* and *oqxAB*) were low (≤122 counts per million (cpm) mapped reads) (Figure 2). The remaining resistance genes were located on plasmids. Resistance genes exhibiting high levels of expression (300 cpm) were apparent in 1_GR_13 (*blaTEM-1B, blaVIM-27, sul1, aph(3’)-Ia*), 2_GR_12 (*aac(6’)-Ib, catA1, blaKPC-2*), 16_GR_13 (*aac(6’)Ib-cr, aac(3)-IIa, blaCTX-M-15, blaTEM-1B, blaOXA-48*) and 20_GR_12 (*blaKPC-2, aac(6’)Ib*). Counts for *aac(6’)-1b* and *aac(6’)-1b-cr* in 2_GR_12 and 20_GR_12 were grouped. The gene *aac(6’)-1b-cr* is a shortened version of *aac(6’)-1b* and both were identified in the same genome position, hence, only *aac(6’)-1b* is displayed in Figure 2. Relative expression did not differ significantly when analysing passed reads alone or all reads. All highly expressed genes were detected within 6 hours as per the real-time detection emulation. As anticipated, low levels of expression were observed for fosfomycin (*fosA*), tetracycline (*tet(A), tet(B)*) and macrolide (*mph(A)*) resistance.

**Figure 2.**
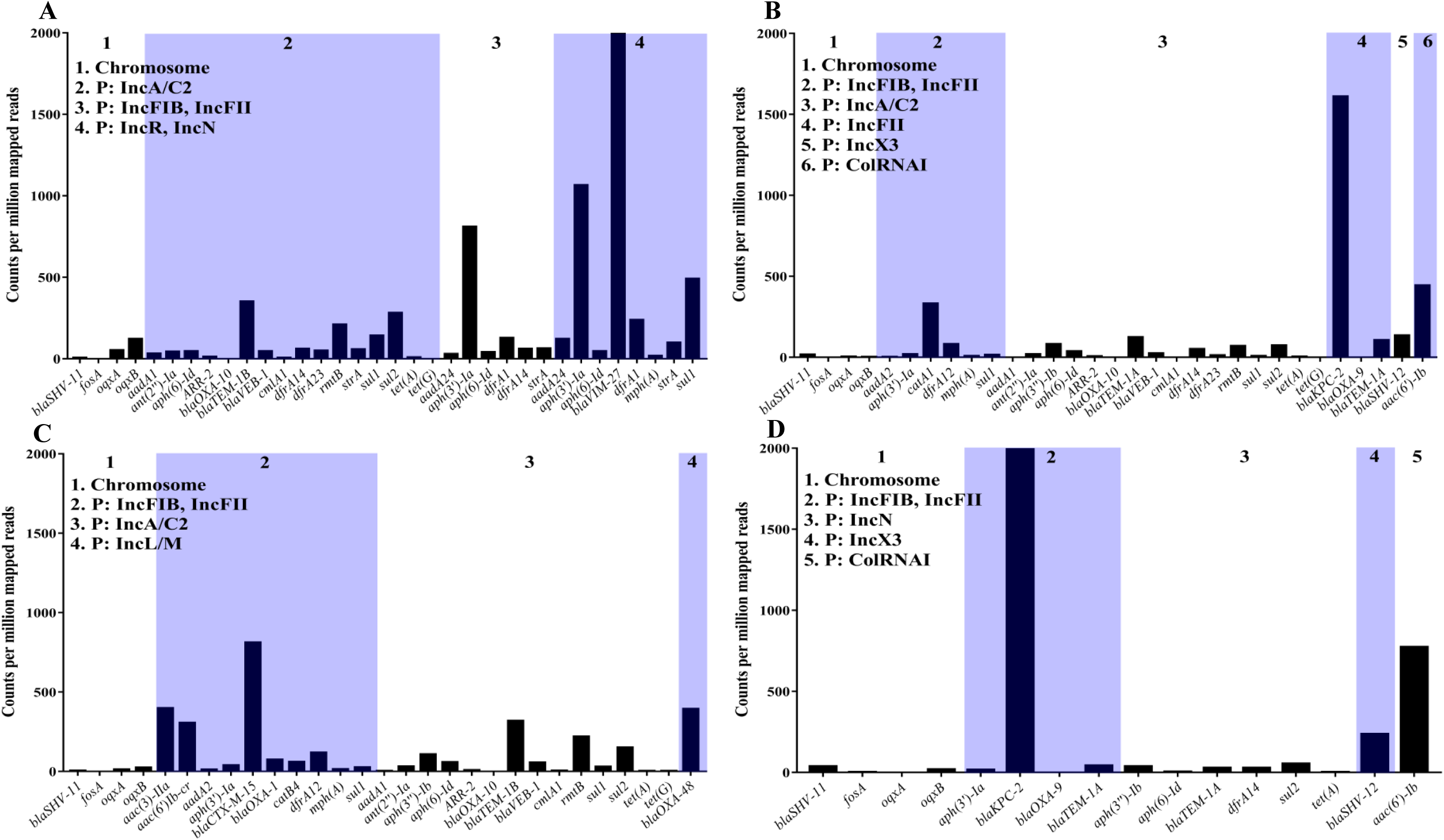
Expression of resistance genes determined using counts per million mapped reads. Due to differing levels of rRNA depletions across samples, reads mapping to rRNA were removed. Strains investigated include (**A**) 1_GR_13, (**B**) 2_GR_12, (**C**) 16_GR_13 and (**D**) 20_GR_12. X-axis depicts the resistance genes and are grouped based on the location in the genome where P indicates a plasmid followed by replicon identity. Albacore 2.2.7 base-called sequences were used and all reads (pass and fail) were included in this analysis.

A subset of 11 resistance genes which represent resistance across various classes of antibiotics were investigated to validate differential gene expression in these RNA extractions via qRT-PCR. These included resistance towards aminoglycosides (*aac(6’)Ib, strA*), β-lactams (*blaKPC-2, blaOXA-10, blaTEM-1*), phenicols (*cmlA1*), trimethoprim (*dfrA14*), fosfomycin (*fosA*), quinolone (*oqxA*), sulphonamides (*sul2*) and tetracyclines (*tet(A)*). A similar trend was observed between direct RNA sequencing and qRT-PCR results (Spearman’s rank correlation coefficient: 0.83; Pearson correlation: 0.86) (Figure 3). The highest expression of a resistance gene was observed for *blaKPC-2* although only one copy was present in a lower copy number plasmid in 2_GR_12 and 20_GR_12 (Figure 2, Figure 3 and Table 1). Additionally, low levels of expression for *fosA* and *tet(A)* were apparent despite exhibiting resistance towards fosfomycin and tetracycline (Figure 3, Supplementary Table S1). Direct RNA sequencing was unable to detect low levels of expression whilst qRT-PCR could detect these genes (Figure 3).

**Figure 3.**
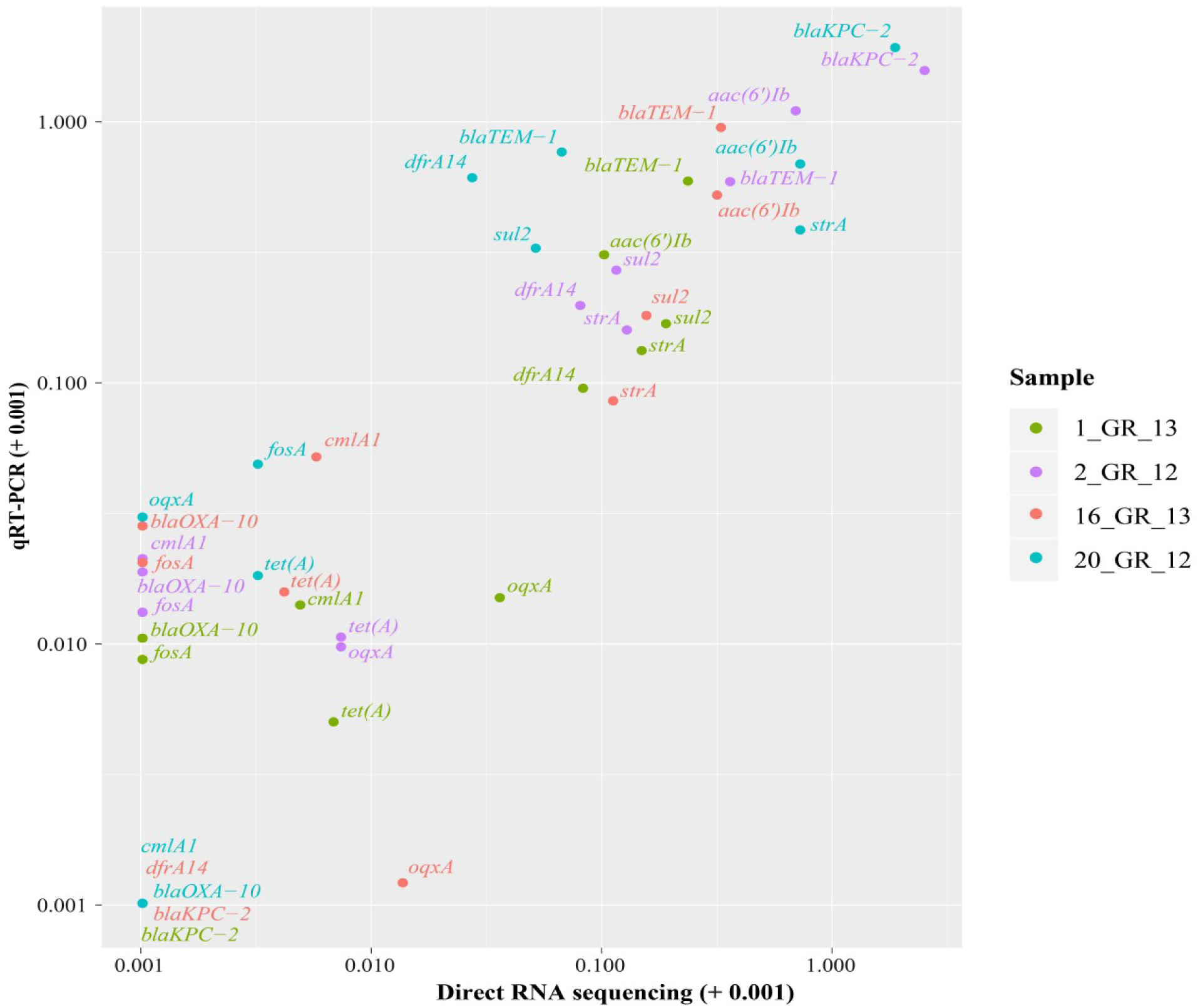
Correlation between resistance genes detected via direct RNA sequencing and validated using qRT-PCR. Relative expression was calculated via normalizing to the housekeeping gene, *rpsL* for both direct RNA sequencing (log2(gene/*rpsL*) and qRT-PCT (2^*-ΔΔCT*^). Due to high similarity between certain genes, several primers recognise more than one gene. These include *aac(6’)Ib: aac(6’)Ib-cr, aadA24*; *strA: aph(3’’)-Ib* and *blaTEM-1: blaTEM-1A, blaTEM-1B.*

Across the transcriptome, antibiotic resistance genes were identified to harbour high differential expression between isolates (Figure 4). Virulence genes were comparable across these strains similar to all remaining or background genes. The top differentially expressed genes were determined (Supplementary Figure S5) and several were associated with polymyxin resistance pathways. Heightened expression was seen in polymyxin-resistant isolates 1_GR_13, 2_GR_12, 16_GR_13 in comparison to the single susceptible isolate in particular, genes associated with Ara4N synthesis. These genes include 4-deoxy-4-formamido-L-arabinose-phosphoundecaprenol deformylase (ArnD), UDP-4-amino-4-deoxy-L-arabinose formyltransferase and UDP-4-amino-4-deoxy-L-arabinose-oxoglutarate aminotransferase.

**Figure 4.**
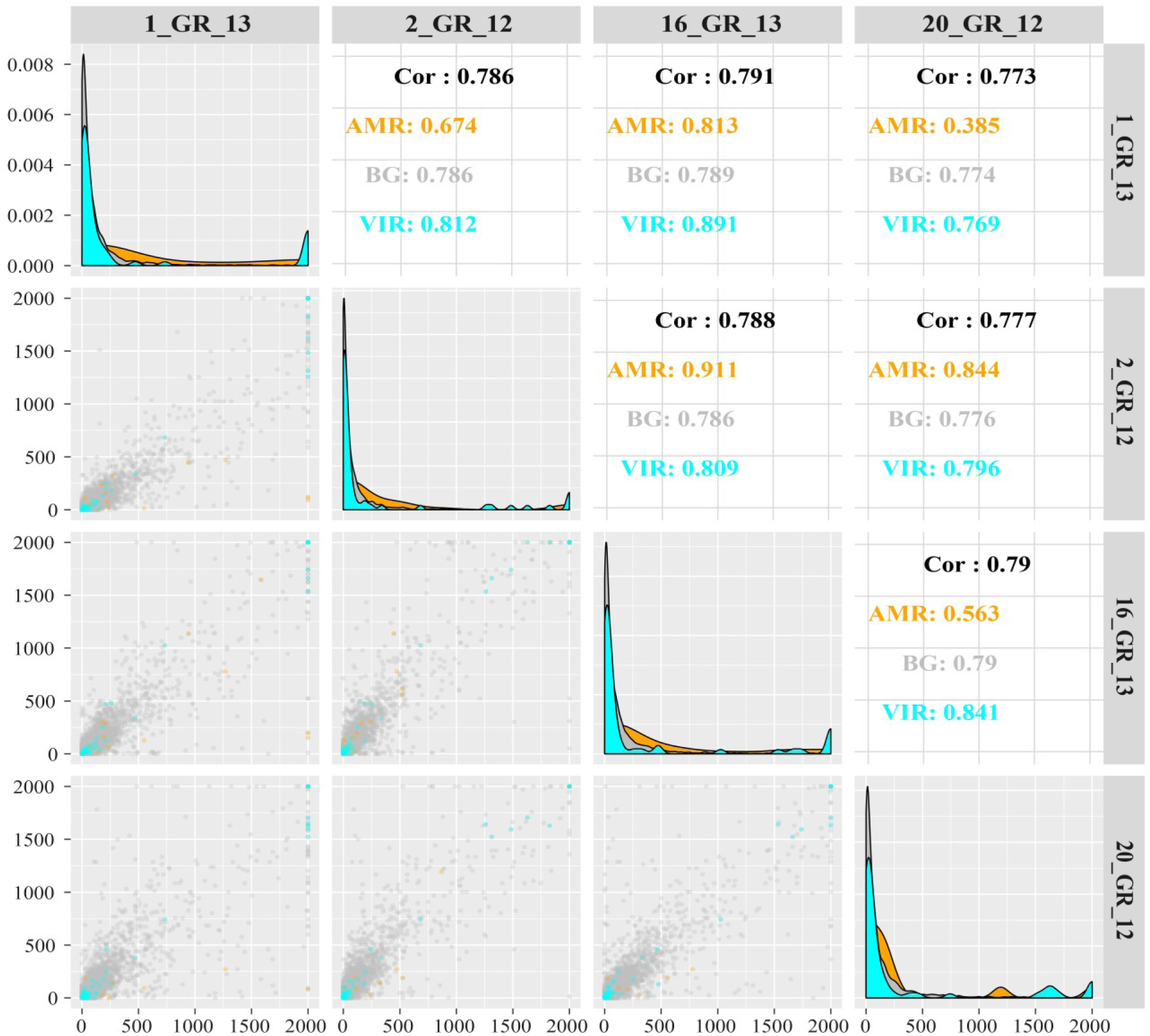
Correlation between gene expression in counts per million mapped reads (capped at 2000 cpm) observed in direct RNA sequencing between the four XDR *K. pneumoniae* isolates. Top panels display spearman correlation coefficients. The diagonal panel displays the histogram of gene expression levels (in cpm) for each sample. Colours indicate categorization of gene: antimicrobial resistance genes (AMR) (as per ResFinder 3.0), virulence genes (VIR) (determined via RAST) and all other genes or background genes (BG) are displayed.

### Transcriptional biomarkers for polymyxin resistance

Three of the isolates harboured resistance towards polymyxins via disruptions in *mgrB* which included 1_GR_13, 2_GR_12 and 16_GR_13. 1_GR_13. These isolates have an insertion sequence (IS) element, IS*Kpn26*-like, at nucleotide position 75 in the same orientation as *mgrB*. 2_GR_12 also contained an insertion at the same position, however, in the opposite orientation and additional mutations in *phoP (*A95S) and *phoQ (*N253T). 16_GR_13 possessed an IS element, IS*1R*-like, 19 bp upstream of *mgrB*. Direct RNA sequencing revealed only low level expression of *mgrB* in isolates (1_GR_13 (78.4 cpm), 2_GR_12 (16.3 cpm), 16_GR_13 (0 cpm), 20_GR_12 (2.3 cpm)). The expression levels of various genes associated with this pathway were verified via qRT-PCR which include genes *phoP, phoQ, pmrA, pmrB, pmrC, pmrD, pmrE, pmrH* and *pmrK (*Figure 5). Direct RNA sequencing revealed a slight increase in transcription of *phoPQ (*≥2-fold) relative to the expression in 20_GR_12. A ≥13-fold increase in expression was observed for *pmrH* and ≥8-fold elevation for *pmrK.* Similar trends for expression were also reported using qRT-PCR (Figure 5B).

**Figure 5.**
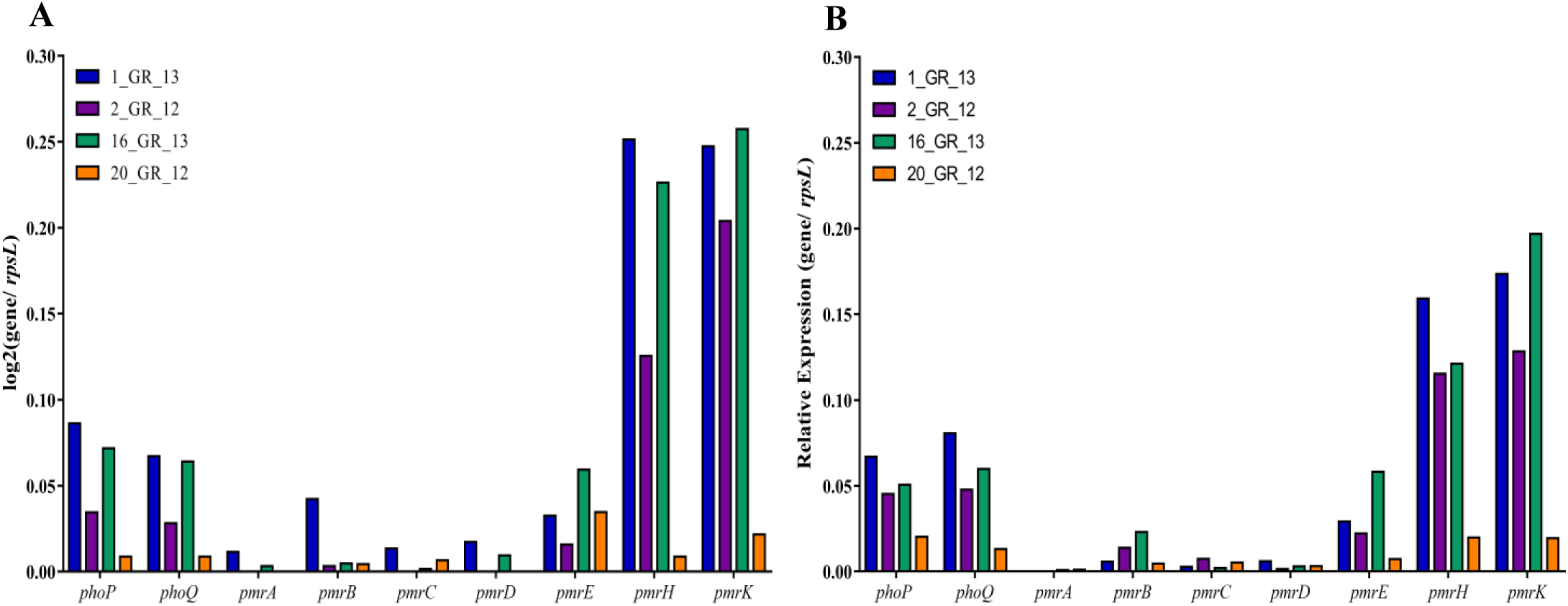
Expression of genes associated with the polymyxin resistance pathway. Comparison between (**A**) direct RNA sequencing and (**B**) qRT-PCR. Isolates harbouring resistance to polymyxins (MIC: >2 µg/mL) include 1_GR_13, 2_GR_12 and 16_GR_13.

## DISCUSSION

XDR *K. pneumoniae* pose as a major threat to modern medicine with rapid diagnostics critical to discern appropriate treatment options (1,6). The MinION sequencing technology employed in this study has great potential to detect antibiotic resistance in a timely manner, as shown with four XDR *K. pneumoniae* isolates. This method was able to resolve both the assembly of plasmids harbouring high levels of resistance (through DNA sequencing) and the expression from the resistome in the absence of antibiotic treatment (through RNA sequencing).

The ability for ONT to sequence long fragments of DNA has significantly aided the assembly of bacterial genomes and plasmids (16-18). In this study, multiple megaplasmids (≥100 kbp) were identified which were previously unresolved via Illumina sequencing (24). These harboured replicons IncA/C2 or a dual replicon, IncFIIK and IncFIB. The IncA/C, IncF and IncN plasmids have been commonly associated with multidrug resistance (46). Although several plasmids in this study revealed similarity to previously reported isolates via NCBI, various sequences deviated. In particular, the IncA/C2 plasmid exhibited multiple regions unique to these isolates. Several IncA/C2 megaplasmids have been previously described which harbour various resistance genes, however, the extent of resistance in our study has yet to be unveiled (47,48). Prior studies have shown the IncFIIK and IncFIB replicons to localise on the same plasmid and also megaplasmids with multidrug resistance (6). The IncFIBpQil plasmid in this study contained various β-lactam resistance genes (*blaKPC-2, blaOXA-9, blaTEM-1A*) which has been identified previously (49). Similarly, *blaOXA-48* segregated with the IncL/M replicon (50,51), however, deviations in this plasmid were identified.

The real-time analysis capability entailed in MinION sequencing has the potential to rapidly determine the antibiotic resistance profile. Previously, this device has been utilised to rapidly assemble bacterial genomes, discern species and detect antibiotic resistance (12-15). This study investigated the potential time required to discern resistance via a real-time emulation as previously described (17). The majority (≥70%) of resistance genes were detected via DNA sequencing within 2 hours. However, several genes that were not identified in the final assembly were apparent after 2 hours. This may be attributed to the high similarity (≥80%) amongst various genes, in particular, those associated with aminoglycoside, β-lactam, rifampicin and phenicol resistance. Furthermore, the error rate associated with ONT sequencing and the accumulation of these errors over time may result in the false detection of these genes. Nanopore DNA sequencing currently has an accuracy ranging from 80 to 90% which limits its ability to detect mutations (17). Various resistance genes only differ by a few nucleotides which significantly impacts the resistance phenotype and the antibiotics which can be utilised to treat these infections. Furthermore, direct RNA sequencing has an average error rate of 12% (21). Hence, it is essential for the technology to increase its accuracy in order to correctly and rapidly diagnose antibiotic resistance.

Investigating the transcriptome of these isolates can potentially elucidate the correlation between genotype and the subsequent resistant phenotype. One of the advantages of RNA sequencing is that it can identify conditions in which a resistance gene is present but not expressed, potentially resulting in a susceptible phenotype. However, if expression is only induced in the presence of an antibiotic, the absence of RNA transcripts may falsely suggest susceptibility. Direct RNA sequencing revealed high levels of transcription from genes associated with aminoglycoside, β-lactam, sulphonamide and trimethoprim resistance within 6 hours. The detection of quinolone, rifampicin, and phenicol resistance correlated to the levels of transcription within samples. All isolates exhibited low levels of expression for fosfomycin, macrolide and tetracycline resistance, despite exhibiting phenotypic resistance to fosfomycin and tetracycline. Whether this transcription is due to prior exposure to these antibiotics in the clinic and the longevity of this expression post exposure warrants further investigation. The changes in transcription levels in response to antibiotic exposure also need to be assessed in future experiments. Furthermore, the time required to detect resistance may be hindered by the slower translocation speed associated with direct RNA sequencing (70 bases/ second) compared to DNA sequencing (450 bases/ second). Furthermore, insufficient rRNA depletion and low base-calling of data could be impacting the detection of this low level expression.

Another variable to consider when evaluating differential expression is the operon or promoter which can further be explored via cloning. In particular, the highest levels of expression were observed for *blaKPC-2* in 2_GR_12 and 20_GR_12. Alterations in the promoter region have previously been reported to influence high levels of expression (52). Furthermore, despite low levels of transcription for fosfomycin (*fosA*) and tetracycline (*tet(A), tet(G)*), phenotypically these isolates consistently retain resistance (24). FosA, an enzyme involved in the degradation of fosfomycin, is commonly encoded chromosomally in *K. pneumoniae* and a combination of expression and enzymatic activity contributes to resistance (53). Genes *tet(A)* and *tet(G)* encode efflux pumps which, in the absence of tetracycline, are lowly expressed (54). Detecting inducible resistance such as tetracycline resistance highlights one of the advantages of investigating the transcriptome. Additionally, copy number of plasmids can further alter the levels of expression detected for these resistance genes.

In this study we also investigated pathways attributed to polymyxin resistance. Three of these strains exhibited an IS element upstream of within *mgrB*, the negative regulator of PhoPQ (25,26). Elevated expression was apparent for *phoPQ* and also the *pmrHFIJKLM* operon in our polymyxin-resistant isolates harbouring a disruption in *mgrB*. This has previously been witnessed for other *K. pneumoniae* isolates harbouring *mgrB* disruptions and is a potential transcriptional marker for polymyxin resistance (27,43,55,56). However, this study is limited to four isolates and one mechanism associated with polymyxin resistance. Other pathways have previously been identified including the role of other TCSs such as CrrAB (57). The ability to use relative expression of key genes to detect polymyxin resistance requires further validation, including an increased sample size of resistant and non-resistant isolates. Furthermore, additional functional experiments such as complementation assays would be required in order to validate the contribution of a certain mutation to the transcriptome and subsequent resistance.

This study has utilised MinION sequencing to assemble four XDR *K. pneumoniae* genomes and has revealed several unique plasmids harbouring multidrug resistance. The vast majority of this resistance was detectable within 2 hours of sequencing, though a number of resistance genes were identified that were not present in the final assembly. Exploiting this analysis in real-time would allow for a rapid diagnostic, however, the presence of a resistance gene does not necessarily indicate resistance is conferred and requires additional phenotypic characterisation. This research also established a methodology and analysis for bacterial direct RNA sequencing. The differential expression of resistance genes were successfully detected via this technology and can be exploited for bacterial transcriptomics. Once base-calling algorithms have been optimised, this could allow for a whole transcriptome interrogation of full length transcripts regulated by operons, where more than one gene is co-expressed in a transcript, and the evaluation of the poorly characterised epitranscriptome. This research established a methodology and analysis for bacterial direct RNA sequencing. The differential expression of resistance genes were successfully detected via this technology and can be exploited for bacterial transcriptomics. Overall, this study has begun to unravel the association between genotype, transcription and subsequent resistant phenotype in these XDR *K. pneumoniae* clinical isolates, establishing the groundwork for developing a diagnostic that can rapidly determine bacterial resistance profiles.

## Supporting information

## ACKNOWLEDGEMENTS

We would like to acknowledge Dr Ilias Karaiskos and Dr Helen Giamarellou for providing the bacterial strains in this study. We also acknowledge Dr Evangelos Bellos for his guidance on the RNA sequencing analysis and Dr Devika Ganesamoorthy for the initial advice on the direct RNA sequencing library preparation.

