## Supplementary material for "Evaluating the Genome and Resistome of Extensively Drug-Resistant *Klebsiella pneumoniae* using Native DNA and RNA Nanopore Sequencing"

**SUPPLEMENTAL MATERIAL**

This pdf contains:

Table S1-5

Figures S1-5

References

**Table S1.** Minimum inhibitory concentrations of the four *Klebsiella pneumoniae* clinical isolates

| Strain[a] | Resistance Profile[b] | | | | | | | | | | | | | | | | | | | | | | | |
|---|---|---|---|---|---|---|---|---|---|---|---|---|---|---|---|---|---|---|---|---|---|---|---|---|
| | 1 | | | 2 | 3 | 4 | | 5 | 6 | | | 7 | 8 | 9 | 10 | 11 | 12 | 13 | 14 | 15 | 16 | | 17 | |
| | AMK | GEN | TOB | CPT | TZP | IPM | MEM | CFZ | FEP | CTX | CAZ | FOX | CIP | SXT | TGC | ATM | AMP | SAM | CHL | FOF | CST | PMB | MIN | TET |
| 1_GR_13 | >64 | >64 | >64 | >32 | >128/ >64 | ≥64 | >32 | >64 | >64 | >32 | >64 | >64 | >16 | >64/ 128 | ≥2 | >64 | >64 | >64/ >64 | 64 | ≤19 | 16 | ≥8 | ≥16 | >64 |
| 2_GR_12 | >64 | >64 | >64 | >32 | >128/ >64 | ≥64 | >32 | >64 | >64 | >32 | >64 | >64 | >16 | >64/ 128 | 2 | >64 | >64 | >64/ >64 | >64 | ≤20 | 16 | 8 | 16 | >64 |
| 16_GR_13 | >64 | >64 | >64 | >32 | >128/ >64 | ≥8 | ≥16 | >64 | >64 | >32 | >64 | 64 | >16 | >64/ 128 | ≥2 | >64 | >64 | >64/ >64 | 32 | ≤19 | 64 | 64 | ≤8 | >64 |
| 20_GR_12 | 64 | 1 | 64 | >32 | >128/ >64 | ≥8 | ≥8 | >64 | ≥16 | >32 | >64 | 64 | >16 | >64/ 128 | ≥2 | >64 | >64 | >64/ ≥64 | ≥32 | ≤22 | 0.125 | ≤0.25 | 16 | >64 |

[a]Strain identification, numerical order catalogued at IMB_Country (GR:Greece, BR:Brazil)_last two digits of isolation year.

[b]Antibiotic resistance as determined by broth microdilution according to CLSI guidelines except fosfomycin (disk diffusion) and tigecycline which followed EUCAST breakpoints. Antibiotic classes tested include **1**, Aminoglycosides (Amikacin, AMK; Gentamicin, GEN; Tobramycin, TOB); **2**, Anti-MRSA cephalosporins (Ceftaroline, CPT); **3**, Antipseudomonal penicillins + β-lactamase inhibitors (Piperacillin-tazobactam, TZP); **4**, Carbapenems (Imipenem, IPM; Meropenem, MEM); **5**, Non-extended spectrum cephalosporins (1$^{st}$ and 2$^{nd}$ generation) (Cefazolin, CFZ); **6**, Extended-spectrum cephalosporins (3$^{rd}$ and 4$^{th}$ generation) (Cefepime, FEP; Cefotaxime, CTX, Ceftazidime, CAZ); **7**, Cephamycins (Cefoxitin, FOX); **8**, Fluoroquinolones (Ciprofloxacin, CIP); **9**, Folate pathway inhibitors (Trimethoprim-sulfamethoxazole, SXT); **10**, Glycylcyclines (Tigecycline, TGC); **11**, Monobactams (Aztreonam, ATM); **12**, Penicillins (Ampicillin, AMP); **13**, Penicillins + β-lactamase inhibitors (Amipicillin-sulbactam, SAM); **14**, Phenicols (Chloramphenicol, CHL); **15**, Phosphonic acids (Fosfomycin, FOF); **16**, Polymyxins (Colistin, CST; Polymyxin B, PMB); **17**, Tetracyclines (Minocycline, MIN; Tetracycline, TET). The two antibiotics for TZP, SXT and SAM were assayed separately and both MICs separated with a /. Shading represents: R, Resistant; I, Intermediate; S, Susceptible. Values indicate MIC in µg/mL except disk diffusion values for fosfomycin in mm.

**Table S2.** Genome assembly comparison

| Strain | Illumina coverage (X) | N50 | Nanopore coverage (X) | N50 | Assembly Method[a] | | | |
|---|---|---|---|---|---|---|---|---|
| | | | | | SPAdes/ npScarf | SPAdes/ Unicycler | Canu | Minimap2/ Miniasm/ Racon |
| 1_GR_13 | 63 | 242938 | 215 | 8712 | **5289533** (IncFIB), **193063** (IncA/C2), **168359** (IncFIB; IncFII), 61039, 55020, **53489** (IncR; IncN), **6457** | **5181675**, **192771** (IncA/C2), 159172 (IncFIB; IncFII), **108879** (IncFIB), **55018**, **53495** (IncR; IncN) | **5073727**, **226704** (IncA/C2), **184820** (IncFIB; IncFII), **136154** (IncFIB), 100186, **82924** (IncR; IncN), 54495 | **5167584**, **192237** (IncA/C2), **168292** (IncFIB; IncFII), **108582** (IncFIB), **53325** (IncR; IncN) |
| 2_GR_12 | 158 | 196706 | 67 | 5251 | 5485776, 457649 (IncFIB; IncA/C2; IncFII), 60365 (IncFII), 42041 (IncX3), 32458, 21300, **13841** (ColRNAI), 12226 | 3743268, 1694231, 175636 (IncA/C2), 152644 (IncFIB; IncFII), 95481 (IncFIB), **43380** (IncX3), 28913, 26127, 16781 (IncFII), 16315, **13841** (ColRNAI) | 5440093, 392198 (IncFIB; IncA/C2; IncFII), 122463 (IncFIB; IncFII), 57534 (IncX3), 26891 (ColRNAI) | 3769033, 1696038, 204124 (IncA/C2), 179356 (IncFIB; IncFII), 158561 (IncFIB), **43085** (IncX3), 2250, 13400 (ColRNAI) |
| 16_GR_13 | 123 | 203529 | 101 | 5012 | **5426917**, **186908** (IncFIB; IncFII), **154971** (IncA/C2), **63588** (IncL/M), 37608, 35578, **5225**, 4426 (ColRNAI), **3703** | **5426765**, **187670** (IncFIB; IncFII), **155161** (IncA/C2), **63589** (IncL/M), **5234**, **4940** (ColRNAI), 2156 (ColRNAI) | **5400611**, **199904** (IncFIB; IncFII), **180542** (IncA/C2), **83467** (IncL/M), 14853, 11623, 9308, 9243, 7568, 7089 | **5410256**, **186950** (IncFIB; IncFII), **154635** (IncA/C2), **63299** (IncL/M), 5100, 4900 |
| 20_GR_12 | 262 | 256217 | 115 | 10151 | **5391578**, **163468** (IncFIB; IncFII), **50940** (IncN), 50856 (IncX3), 12578 (ColRNAI) | **5395894**, **170467** (IncFIB; IncFII), **50979** (IncN), **43380** (IncX3), **13841** (ColRNAI), 4645 | 5342491, **192947** (IncFIB; IncFII), 74844 (IncX3), 72588 (IncN), 25624 (ColRNAI) | **5380057**, **169880** (IncFIB; IncFII), **50636** (IncN), **43157** (IncX3), 13600 (ColRNAI) |

[a] Output of genome assembly identifies circular sequences (**bold**) and in brackets, plasmid replicons present on these contigs.

**Table S3.** Real-time emulation of time to detect resistance genes from DNA and RNA sequencing

| Isolate | Time (mins) | DNA | RNA |
|---|---|---|---|
| | | Resistance gene/s detected[a] | |
| 1_GR_13 | 10 | **oqxA** (Q), **dfrA1** (Tr), **dfrA14** (Tr), **dfrA23** (Tr), **blaVIM-27**[#] (B), **sul1**/3 (S), **strA** (A), **aph(3')-Ia**/c (A), strB (A), aadB (A), **mph(A)** (M), **blaTEM-1B**[#] (B), **oqxB** (Q), **rmtB2** (A) | **sul1**/ 3 (S) |
| | 30 | **sul2** (S), **ARR-2**/3/6 (R), **blaOXA-10**[#] (B), **blaVEB-1**[#] (B), **tet(G)** (T), **cmlA1** (P), floR (P), **blaSHV-11**[#] (B), **fosA** (F) | **aph(3')-Ia/c** (A), **blaTEM-1B**[#](B), **blaVIM-27**[#] (B), **fosA** (F) |
| | 60 | ARR-3 (R), aac(6')Ib[#] (A), ARR-7 (R), aac(2') (A), cml (P) | ARR-7 (R) |
| | 120 | aac(3')-IIIc (A) | **dfrA14** (Tr), **oqxA** (Q) |
| | 300 | aac(3')-IIIb (A), aac(6')-Ic (A), | strB (A), **sul2** (S), **blaVEB-1**[#] (B), **dfrA1** (Tr), **dfrA23** (Tr), **strA** (A), **blaSHV-11**[#] (B) |
| | 600 | blaPAO (B), aph(3')-IIb (A), aph(6)-Ic (A) | aadB (A) |
| | 900 | catpC233 (P), blaOKP (B) | **rmtB2** (A) |
| | 1200 | **tet(A)** (T) | - |
| 2_GR_12 | 10 | **blaKPC-2**[#] (B), **blaTEM-1A**[#](B), **aac6Ib/-cr**[#] (A), **cmlA1** (P), **dfrA12** (Tr), **sul1**/ 3 (S) | **blaKPC-2**[#] (B), **catA1** (P) |
| | 30 | **rmtB2** (A), **oqxA** (Q), **dfrA14** (Tr), blaPAO (B) | **aac6Ib/-cr**[#] (A) |
| | 60 | **oqxB** (Q), strA (A), strB (A), **aph(3')-1a**/c (A), **sul2** (S), **blaSHV-11/12**[#] (B), **mph(A)** (M), **catA1** (P), **tet(G)** (T), **blaOXA-9**[#] (B), **aadA1/2**[#] (A), fusB (Fu), **aac(6')**/aph(2") (A) | **sul2** (S) |
| | 120 | **blaOXA-10**[#] (B), floR (P), **ARR-2/3/6** (R), aadB (A), **fosA** (F), **blaVEB-1**[#] (B), cml (P), **dfrA23** (Tr) | **blaTEM-1A**[#] (B), **blaSHV-11/12**[#](B) , **ARR-2**[#](R), aadB (A) |
| | 300 | aac(3')-IIIc (A), catpC233 (P), aac(3')-IIIb (A), ARR-3 (R) | **rmtB2** (A), **blaVEB-1**[#](B), strB (A), strA (A), **dfrA14** (Tr) |
| | 600 | **tet(A)** (T) | **dfrA12** (Tr) |
| | 900 | aph(6)-Ic (A) | - |
| | 1200 | - | **fosA** (F) |
| 16_GR_13 | 10 | blaOXA-436 (B), **sul1**/3 (S), **dfrA12** (Tr), **fosA** (F), aadB (A), strA (A), **sul2** (S), strB (A), blaCTX-M-64 (B) | - |
| | 30 | **blaOXA-48**[#] (B), **aph(3')-1a**/c (A), **mph(A)** (M), **rmtB2** (A), floR (P), **blaCTX-M-15**[#] (B), **aac(6')Ib-cr**[#] (A), **blaOXA-1**[#] (B), **oqxB** (Q), **oqxA** (Q), **blaTEM-1B**[#] (B), **blaVEB-1**[#] (B), **cmlA1** (P) | **aac(3')-IIa**[#] (A), **sul2** (S), blaOXA-436 (B) |
| | 60 | **tet(G)** (T), **blaOXA-10**[#] (B), **blaSHV-11**[#] (B), **aac(3')-IIa**[#] (A), **ARR-2/3/6** (R) | **rmtB2** (A), **blaOXA-48**[#] (B), strA (A), **blaTEM-1B**[#] (B) |
| | 120 | **aadA1/2**[#] (A), fusB (Fu), ermT (M), str (A), rmtG (A), **aac(6')**/aph(2") (A) | blaCTX-M-64 (B), **blaCTX-M-15** (B) |
| | 300 | ARR-3 (R), catpC233 (P), cml (P), aph(6)-Ic (A), aac(3')-IIIb (A) | **dfrA12** (Tr), strB (A), **blaVEB-1**[#] (B), **blaOXA-1**[#] (B), **aph(3')-Ia**/c (A), **oqxA** (Q), **sul1**/ 3 (S) |
| | 600 | vanR (V), dfrA14 (Tr) | - |
| | 900 | blaPAO (B) | aadB (A), **aac(6')Ib-cr**[#] (A) |
| | 1200 | aph(3')-IIb (A) | - |
| 20_GR_12 | 10 | **aac(6')Ib/-cr**[#] (A), **blaTEM-1A**[#] (B), **dfrA14** (Tr), **sul2** (S), strB (A), **blaKPC-2**[#] (B), **tet(A)** (T) | **aac(6')Ib/-cr**[#] (A) |
| | 30 | **oqxA** (Q), **blaSHV-11/12**[#] (B), **aph(3')-Ia** (A), **fosA** (F), **blaOXA-9** (B), **oqxB** (Q), aph(6)-Ic (A) | **blaKPC-2**[#] (B) |
| | 60 | - | - |
| | 120 | aac(2') (A), catpC233 (P), aac(3')-IIIb (A) | **dfrA14** (Tr) |
| | 300 | ermT (M), blaPAO (B), **aac(6')**/aph(2") (A), catpC221 (P) | **blaSHV-11/12**[#] (B), **blaTEM-1A**[#] (B) |
| | 600 | rmtf (A), ermG (M) | **sul2** (S) |
| | 900 | aac(3')-IIIc (A), aac(6')-Ic (A) | - |
| | 1200 | vatB (M), ARR-2/3/6 (R), aadB (A) | - |

[a] Resistance genes detected as per the real-time emulation. **Bold** represents gene or gene family detected in final assembly and [#] indicates more than three genes grouped within this family. Genes displayed in order of time detected. An underline identifies genes with a low mapping quality reads (MAPQ: ≤10). Brackets represent class of antibiotic this gene confers resistance which includes: A, aminoglycoside; B, β-lactam; F; fosfomycin; Fu; fusidic acid, M, macrolide; P, phenicol; Q, quinolone; R, rifampicin; S, sulphonamide; T, tetracycline; Tr, trimethoprim; V, vancomycin.

**Table S4.** Oligonucleotides used in this study for qRT-PCR

| Assay | Gene | Forward sequence (5' to 3') | Reverse sequence (5' to 3') | Reference[a] |
|---|---|---|---|---|
| Acquired resistance | | | | |
| | *aac(6')Ib*[#] | TTG CAA TGC TGA ATG GAG AG | TGG TCT ATT CCG CGT ACT CC | TS |
| | *blaKPC-2* | TGG CTA AAG GGA AAC ACG AC | TAG TCA TTT GCC GTG CCA TA | TS |
| | *blaOXA-10* | GGT GGG TTG AGA AGG AGA CA | ATG ATT TTG GTG GGA ATG GA | TS |
| | *blaTEM-1*[#] | AAG CCA TAC CAA ACG ACG AG | TTG CCG GGA AGC TAG AGT AA | TS |
| | *cmlA1* | AAT GGG ATG CCT GAT AGC TG | ACC CAC TAG CCA CAT TGG AG | TS |
| | *dfrA14* | TTT GAA TCT ATG GGC GCA CT | ATG GCC TCT TCG ATT GAC TG | TS |
| | *fosA* | CGT GGC GTT TTA TCA GCA G | ACA GGC ACA GCC ACA AAT C | TS |
| | *oqxA* | GCG ATG ATG CTC TCC TTT CT | GAT CGA CTT CAC CAG CAC CT | TS |
| | *strA*[#] | ACT CTT CAA TGC ACG GGT CT | CCA GTT CTC TTC GGC GTT AG | TS |
| | *sul2* | GAT ATT CGC GGT TTT CCA GA | GTC TTG CAC CGA ATG CAT AA | TS |
| | *tetA* | TTG GCA TTC TGC ATT CAC TC | GAA GGC AAG CAG GAT GTA GC | TS |
| Polymyxin resistance | | | | |
| | *phoP* | ATT GAA GAG GTT GCC GCC CGC | GCT TGA TCG GCT GGT CAT TCA CC | TS |
| | *phoQ* | GCA TAT CTT CCC GCT GTC AT | GCT AAC GCT ATA GCC CAC CA | TS |
| | *pmrA* | GAT GAA GAC GGG CTG CAT TT | ACC GCT AAT GCG ATC CTC AA | 1 |
| | *pmrB* | TGC CAG CTG ATA AGC GTC TT | TTC TGG TTG TTG TGC CCT TC | 1 |
| | *pmrC* | GCG TGA TGA ATA TCC TCA CCA | CAC GCC AAA GTT CCA GAT GA | 1 |
| | *pmrD* | GAT CGC AGA GAT TGA AGC CT | GCG TTG CGA ATC TTC AAA GT | 1 |
| | | | GCG TTG CGG ATC TTC AAA GT[1, 16] | TS |
| | *pmrE* | GGG TTG ATC TCT GTG ACA TC | GCC TAC CGT AAT GCC GAC TA | 1 |
| | | GGG TTA ATC TCC GCA ACA TC[16] | TGC ATA TCG CAA TGC TGA CTA[16] | TS |
| | | | GCA TAC CGT AAT GCC GAC TA[1] | TS |
| | *pmrH* | CCG CAT CCG TAG CCT GAA | CGT GGG TCT GGC GAT CAT | 2 |
| | *pmrK* | AGT ATC GGT CAG TGG CTG TT | CCG CTT ATC ACG AAA GAT CC | 1 |
| | *rpsL* | CCG TGG CGG TCG TGT TAA AGA | GCC GTA CTT GGA GCG AGC CTG | 3 |

[a] A (TS) indicates primers were designed in this study. Superscript after a primer designates alternative coding for: [1], 1_GR_13 (ST147) and [16], 16_GR_13 (ST11). [#] Primer recognises multiple genes, *aac(6')Ib: aac(6')Ib-cr, aadA24*, *strA: aph(3'')-Ib* and *blaTEM-1: blaTEM-1A, blaTEM-1B.*

**Table S5.** Highest similarity observed for final assembly contigs when aligned to NCBI database

| Isolate | ST | Contig | Length (bp) | Contig ID | Accession number | Query Coverage | Identity |
|---|---|---|---|---|---|---|---|
| 1_GR_13 | 147 | 1 | **5181675** | C | CP018719.1[*] | 99 | 99 |
| | | 2 | **192771** | P: IncA/C2 | CP008824.1[*] | 76 | 99 |
| | | 3 | **168873** | P: IncFIB$_{pKpn3}$, IncFII$_{pKP91}$ | CP006657.1[*] | 89 | 99 |
| | | 4 | **108879** | P: IncFIB$_{pKPHS1}$ | CP023929.1[*] | 92 | 99 |
| | | 5 | **55018** | - | CP023927.1[*] | 100 | 99 |
| | | 6 | **53495** | P: IncR, IncN | CP023926.1[*] | 92 | 99 |
| 2_GR_12 | 258 | 1 | **5466424** | C | LT216436.1[*] | 99 | 99 |
| | | 2 | 197872 | P: IncFIB$_{pKpn3}$, IncFIIK | CP021713.1[*] | 97 | 99 |
| | | 3 | 175636 | P: IncA/C2 | LT882698.1[*] | 80 | 99 |
| | | 4 | 95481 | P: IncFIB$_{pQil}$ | CP023928.1 | 100 | 99 |
| | | 5 | **43380** | P: IncX3 | CP009776.1[*] | 100 | 99 |
| | | 6 | **13841** | P: ColRNAI | CP010362.1[*] | 99 | 99 |
| 16_GR_13 | 11 | 1 | **5426917** | C | CP018438.1[#] | 97 | 99 |
| | | 2 | **187670** | P: IncFIB$_{pKpn3}$; IncFIIK | CP018693.1[#] | 90 | 100 |
| | | 3 | **155161** | P: IncA/ C2 | KX029331.1[#] | 84 | 99 |
| | | 4 | **63589** | P: IncL/ M$_{pOXA-48}$ | CP018723.1[#] | 100 | 99 |
| | | 5 | **5234** | - | CP025818.1[*] | 99 | 99 |
| | | 6 | **4940** | P: ColRNAI | CP024579.1 | 84 | 99 |
| 20_GR_12 | 258 | 1 | **5395894** | C | CP027160.1[#] | 96 | 99 |
| | | 2 | **170467** | P: IncFIB$_{pKpn3}$; IncFIIK | CP022574.1[#] | 99 | 99 |
| | | 3 | **50979** | P: IncN | CP026053.1[#] | 93 | 99 |
| | | 4 | **43380** | P: IncX3 | CP009776.1[*] | 100 | 99 |
| | | 5 | **13841** | P: ColRNAI | CP010362.1[*] | 100 | 100 |

Query coverage indicates percentage (%) of sequence from NCBI database aligning to contig and identity (%) represents sequence similarity. [#]Indicates the full length of the previously reported sequence is $\geq$90% the length of the contig and [*] is a sequence longer than the contig.

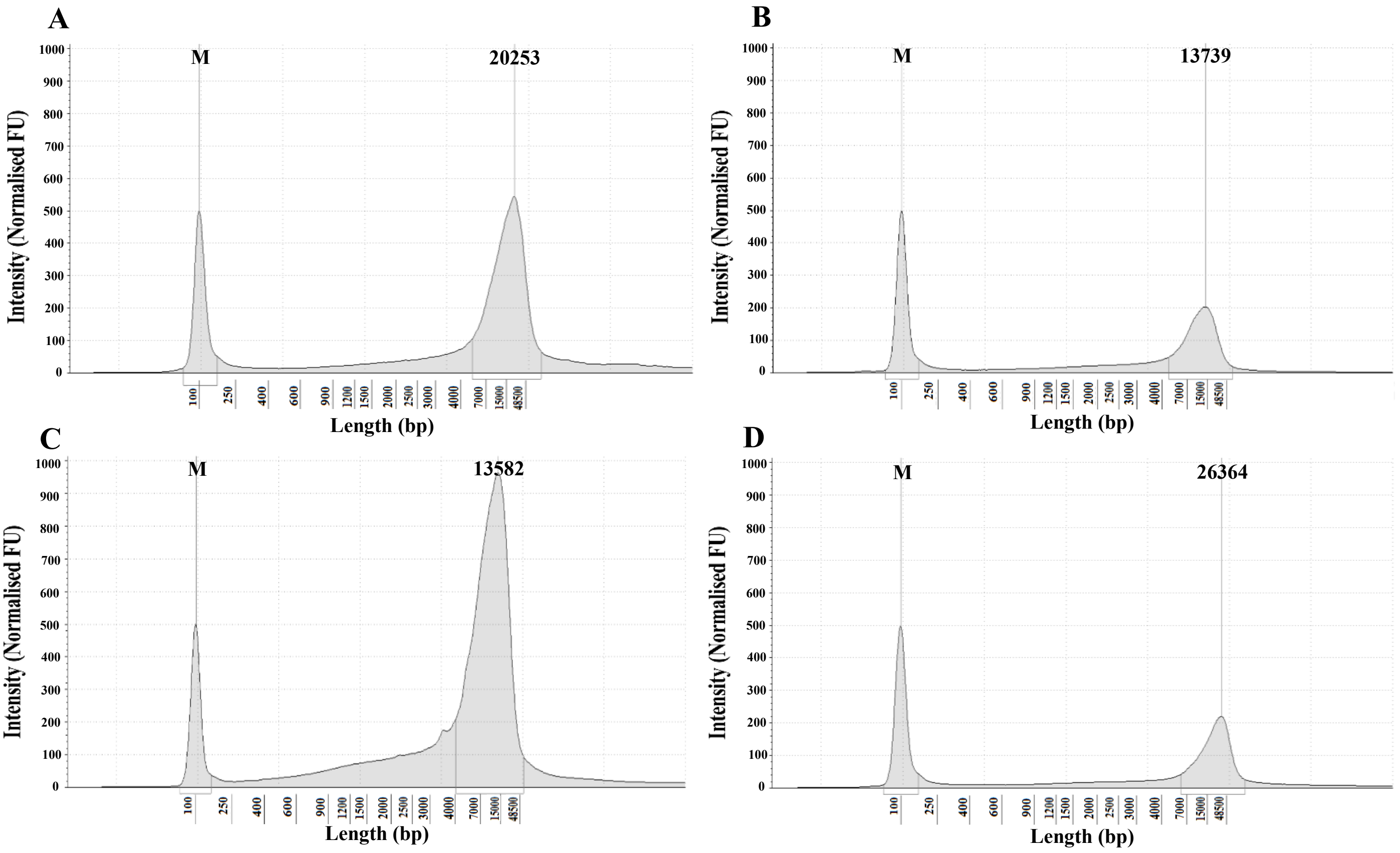


**Figure S1.** Tapestation traces of high molecular weight DNA samples including, (**A**) 1_GR_13, (**B**) 2_GR_12, (**C**) 16_GR_13, (**D**) 20_GR_12. The 100 nt marker is indicated with an M and number indicates the most abundant length.

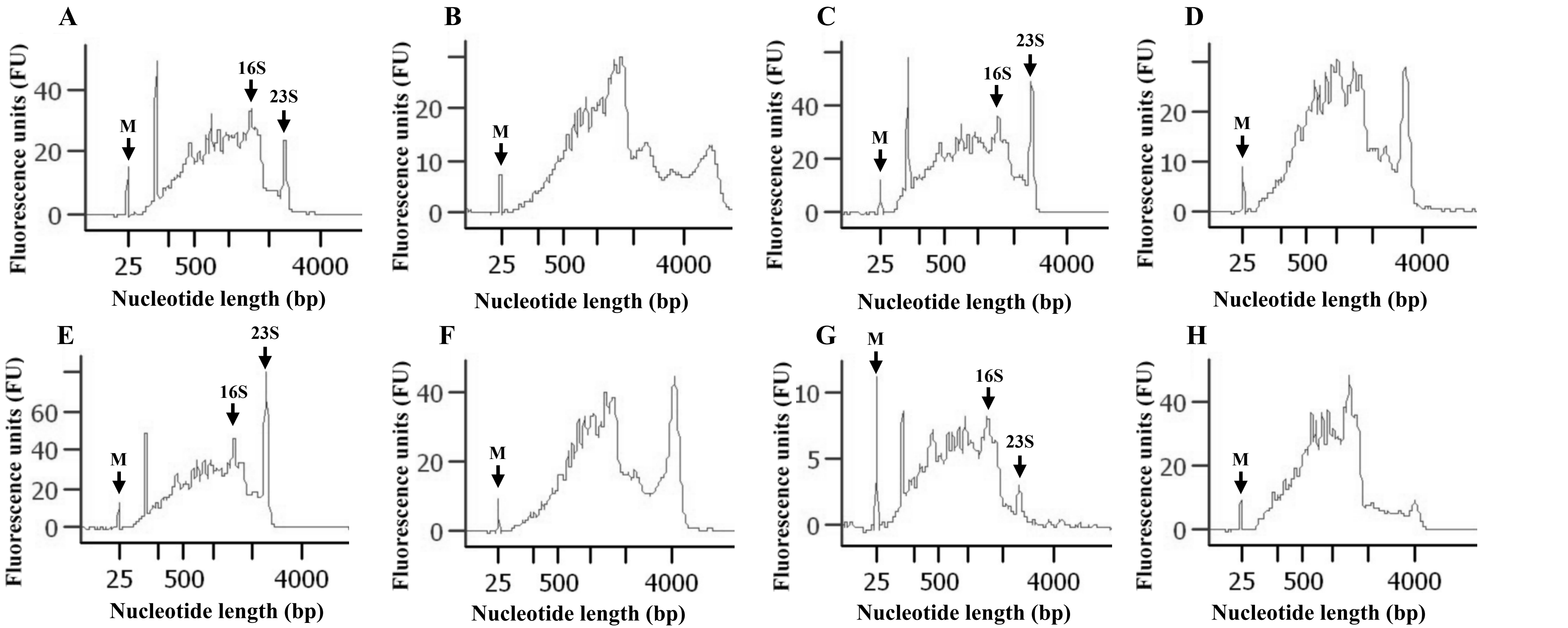


**Figure S2.** Bioanalzyer traces comparing mRNA enrichment and subsequent poly(A) ligation for RNA samples. Sample conditions are as follows: (**A**) 1_GR_13 rRNA depletion, (**B**) 1_GR_13 poly(A) ligation, (**C**) 2_GR_12 rRNA depletion, (**D**) 2_GR_12 poly(A) ligation, (**E**) 16_GR_13 rRNA depletion, (**F**) 16_GR_13 poly(A) ligation, (**G**) 20_GR_12 rRNA depletion, (**H**) 20_GR_12 poly(A) ligation. The 25 nt marker is indicated with an M, presence of 16S rRNA (16S) and 23S rRNA (23S) are also shown.

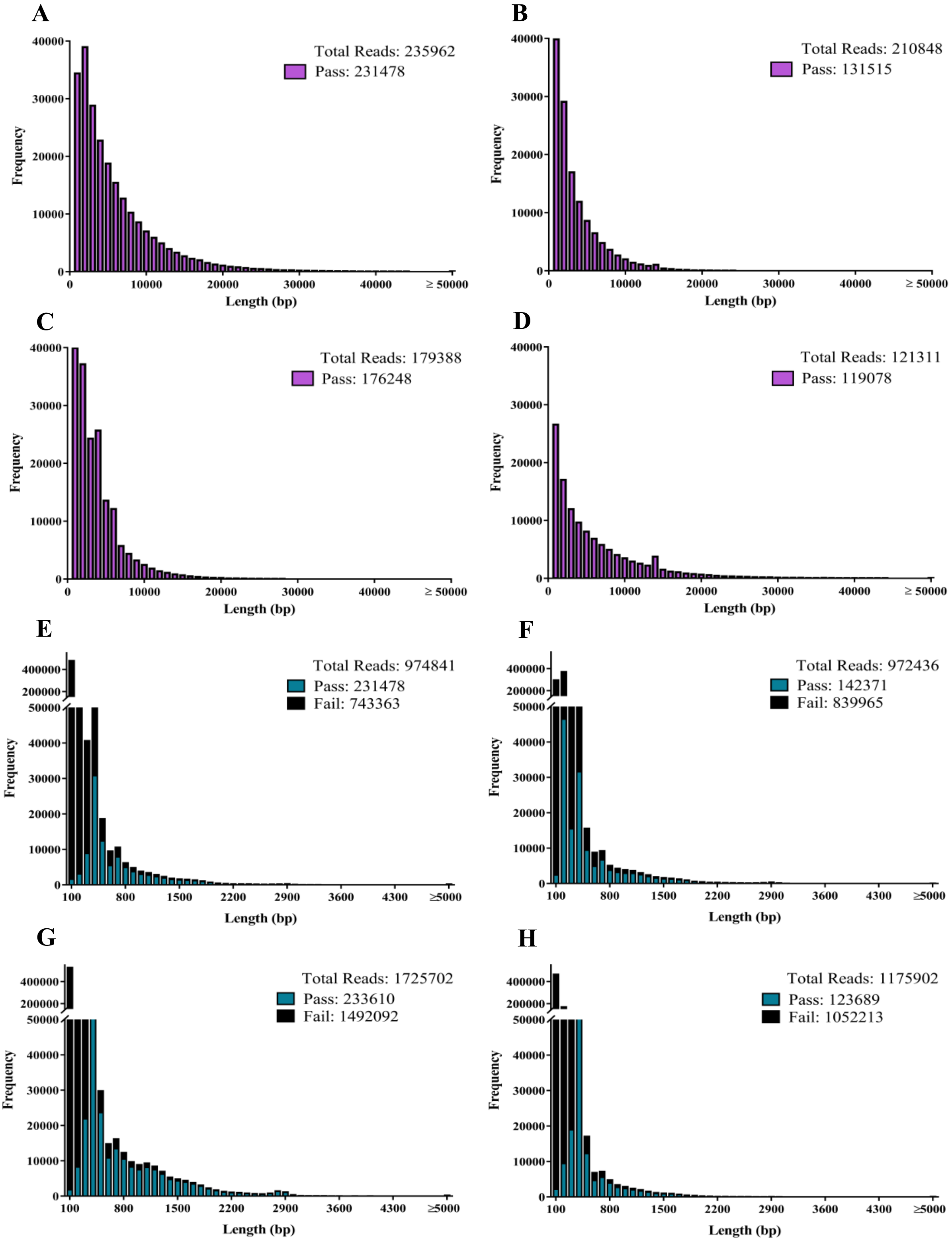


**Figure S3.** Size distribution of reads for sequenced samples. Isolates sequenced by the MinION platform include: (**A**) 1_GR_13 DNA, (**B**) 2_GR_12 DNA, (**C**) 16_GR_13 DNA, (**D**) 20_GR_12, (**E**) 1_GR_13 RNA, (**F**) 2_GR_12 RNA, (**G**) 16_GR_13 RNA, (**H**) 20_GR_12 RNA.

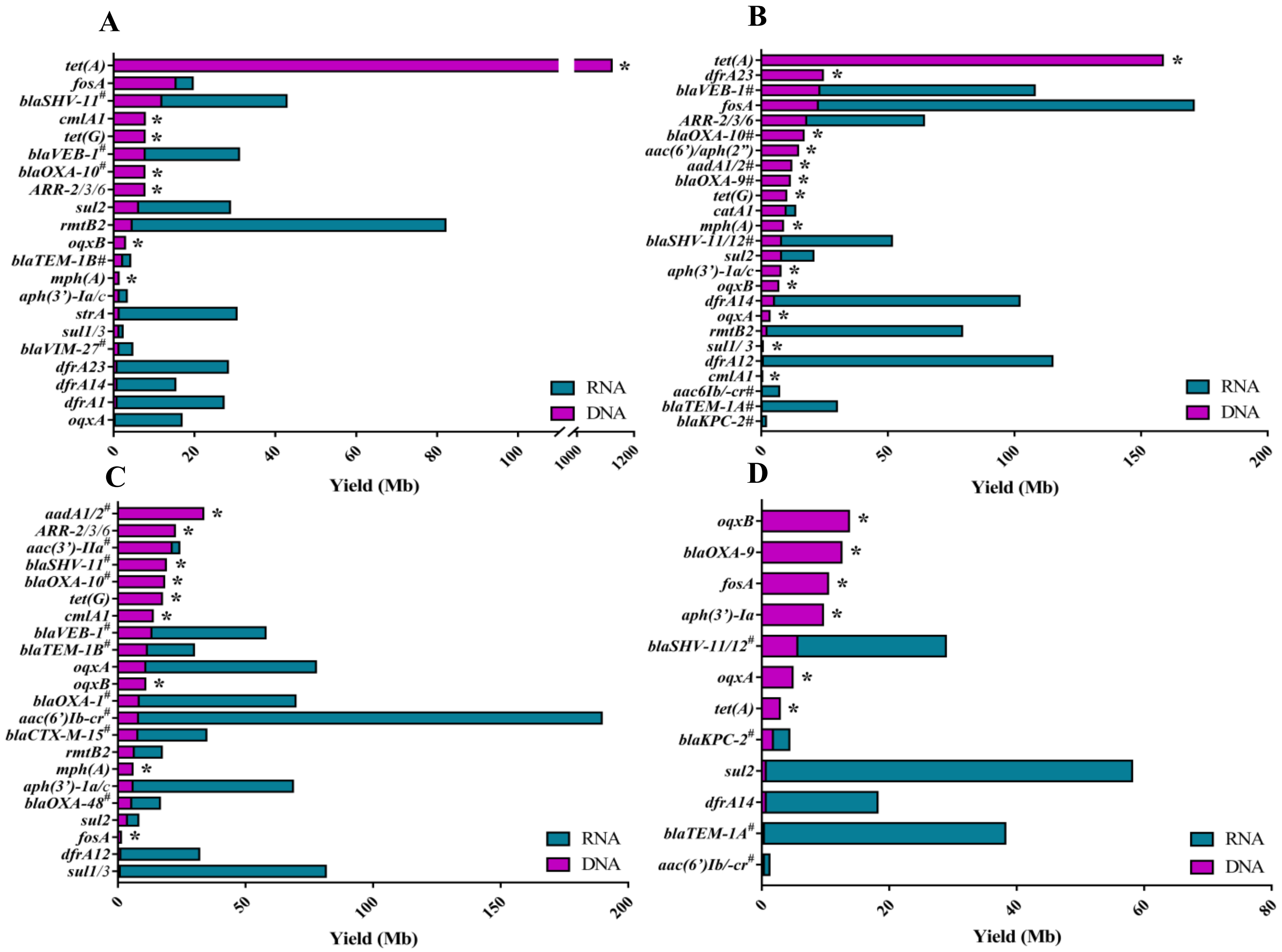


**Figure S4.** Detection of resistance genes via the real-time emulation analysis using DNA or direct RNA MinION sequencing. (**A**) 1_GR_13, (**B**) 2_GR_12, (**C**) 16_GR_13 and (**D**) 20_GR_12. The y-axis displays the resistance genes where an (/) indicates reads detecting more than one gene, (#) is a family of genes (>3) and **bold** displays a gene identified in the final assembly. An asterisk (*) on bars highlights the lack of detection in direct RNA sequencing.

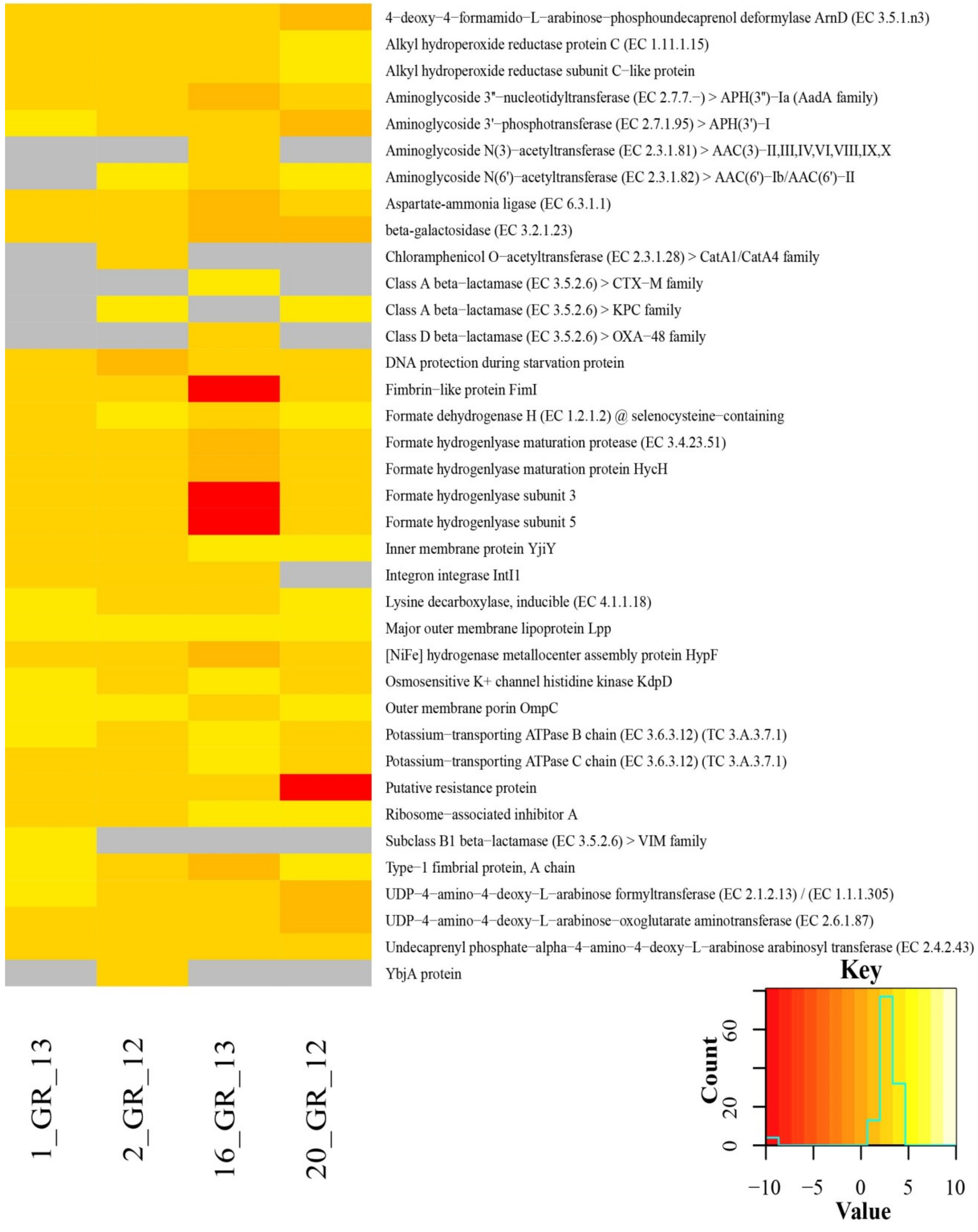


**Figure S5.** Heatmap depicting the top differentially expressed genes across the four *K. pneumoniae* isolates. Expression determined via ONT direct RNA sequencing. Key indicates whether these genes were over-expressed (yellow) or under-expressed (red). Grey indicates the absence of this gene in the isolate.
